# REM negatively predicts statistical learning but not other forms of gist

**DOI:** 10.1101/578492

**Authors:** Nelly Matorina, Jordan Poppenk

**Affiliations:** Department of Psychology, Queen’s University, K7L 3N6; Centre for Neuroscience, Queen’s University, K7L 3N6; School of Computing, Queen’s University, K7L 3N6

**Keywords:** sleep and memory, gist extraction, memory reorganization, gist memory, memory consolidation

## Abstract

Human memory for recent events is believed to undergo reactivation during sleep. This process is thought to be relevant for the consolidation of both individual episodic memories and gist extraction, the formation of generalized memory representations from multiple, related memories. Which kinds of gist are actually enhanced, however, is the subject of less consensus. To address this question, we focused our design on four types of gist: inferential gist (relations extracted across non-contiguous events), statistical learning (regularities extracted from a series), summary gist (a theme abstracted from a temporally contiguous series of items), and category gist (characterization of a stimulus at a higher level in the semantic hierarchy). Sixty-nine participants (30 men, 38 women, and 1 other) completed memory encoding tasks addressing these types of gist and corresponding retrieval tasks the same evening, the morning after, and one week later. Inferential gist was retained over a week, whereas memory for category gist, summary gist, and statistical learning decayed. Higher proportions of REM were associated with worse performance in a statistical learning task controlling for time. Our results support that REM sleep is involved in schema disintegration, which works against participants’ ability to identify regularities within temporal series.

**SIGNIFICANCE STATEMENT:** To gain the most from our experiences, we extract from them the most important elements, or “gist”, with sleep believed to facilitate this process. However, what is referred to as gist varies considerably across studies. We report categorically different mnemonic trajectories of two classes of gist. In particular, we show that gist involving synthesis across relational memories is retained over time, whereas other gists were subject to substantial decay. Moreover, our evidence supports the idea that REM works to discretize, rather than synthesize experiences. Future research should test similar constructs in different tasks to determine whether these findings are generalizable. Our research suggests that patients with reduced REM sleep may experience more interference between similar memories.

## Introduction

To better understand our world, we review the information contents of our experiences for patterns, extracting from those experiences the gist, or essential meaning. ^1^ Fuzzy-trace theory suggests that episodic memory is divided into a detailed memory trace, which preserves the form and characteristics of the information, and a gist memory trace, which preserves the general idea.^1^ There are multiple definitions for gist across different tasks (e.g., a gist representation of semantic associates in a list in the DRM paradigm^2^; information from the generalized series in transitive inference^3^; see Box 1). “Gist extraction” of this kind can take time, and even sleep to emerge: participants who sleep typically demonstrate greater gist memory than those remaining awake for an equivalent period. This pattern has been observed in a variety of common paradigms, including transitive inference,^4^ statistical learning,^5^ and false memory.^6^ One perspective on this gradual emergence of gist is that it reflects the qualitative reorganization of memories during sleep, where new memories emerge that were never learned directly^7^; and several researchers have proposed theoretical frameworks that this occurs either during slow-wave sleep^5^ (SWS) or rapid eye movement (REM) sleep.^8^ However, not all “gists” stand to benefit from such reorganization. For example, remembering the gist of a photograph (such as whether it was taken indoors or outdoors) requires extracting the image’s category without necessarily relating the image to others. We therefore asked the question: do common sleep mechanisms underlie extraction of various gists, and do the various gists benefit differentially?

**Box 1.**
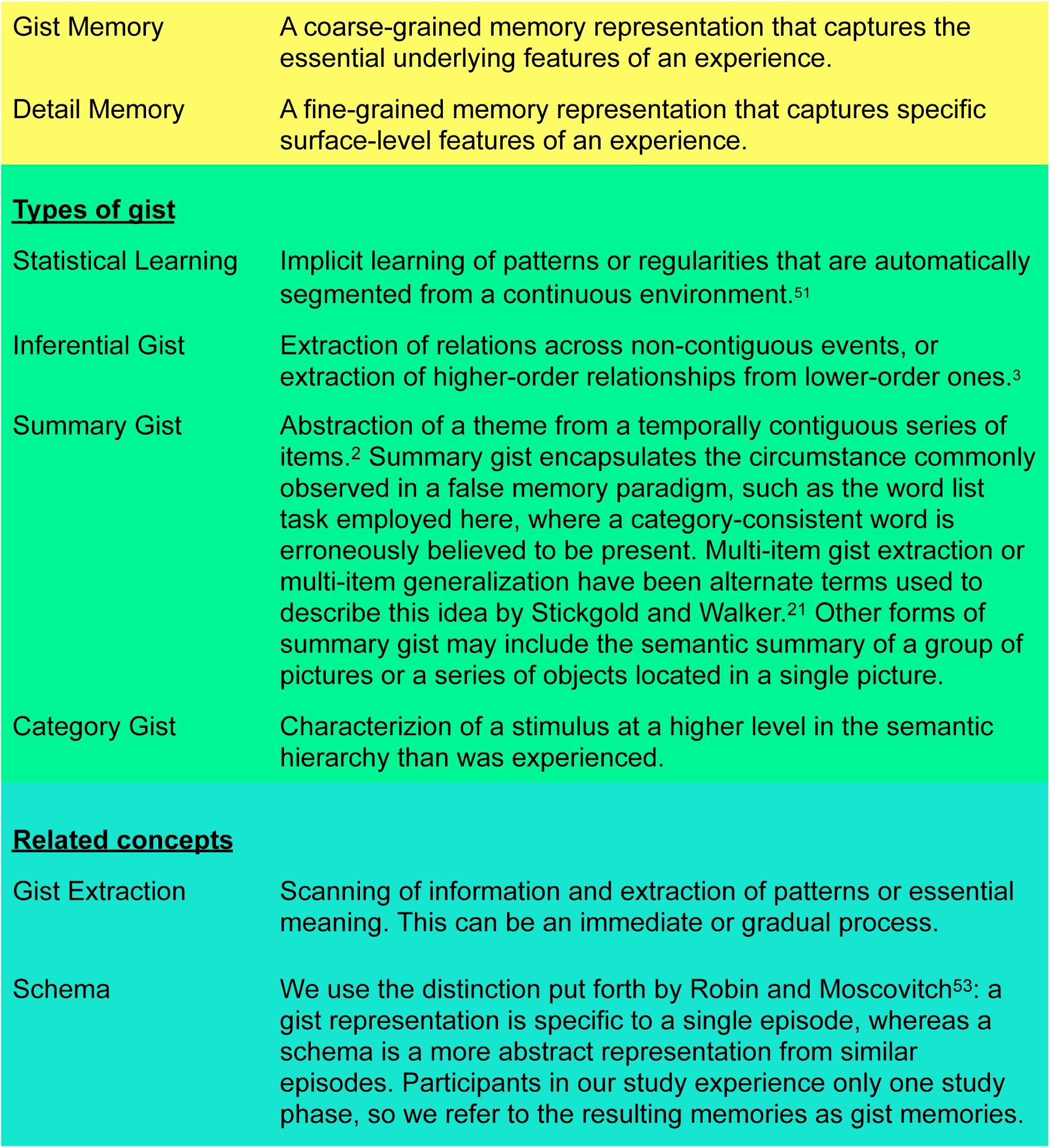
Definitions

During SWS, slow oscillation field potentials (< 1 Hz) are temporally synchronized with thalamo-cortical spindles (7 – 15 Hz) and hippocampal ripples (140-220 Hz).^9^ This process is thought to reactivate memories in the hippocampus, promoting memory consolidation.^10^ Consistent with this “mere reactivation” view, Walker and Stickgold proposed that SWS supports veridical memory consolidation that keeps individual memories distinct.^8^ Under this framework, SWS should preferentially amplify gists internal to a stimulus, such as remembering category gist in the example above. In an alternative model, the information overlap to abstract (IOtA) theory, Lewis and Durrant propose that repeated reactivation of overlapping memory elements leads to strengthening of these shared elements.^11^ Consistent with this theory, duration of SWS correlated with improvement in a statistical learning task^5^ and participants had more slow-wave activity after integrating words into an existing vocabulary.^12^ Under the IOtA account, gist requiring memory integration, such as transitive inference (a task requiring the stitching together of pair relations to make inferences), should preferentially benefit from SWS.

Adding further complexity, Walker and Stickgold proposed cortico-cortical processing in association areas during REM might facilitate associative linking in different cortical areas.^8^ In particular, they proposed REM involvement in three forms of memory integration, all of which contribute to the construction of higher-order schemas: unionization of recent related items, assimilation of new items into established networks, and the abstraction of general rules. Along these lines, higher performance in a probabilistic learning task predicted that more of the subsequent night would be spent in REM sleep,^13^ suggesting that REM is upregulated and promotes reactivation of the same circuits involved in learning.

Here, we took the novel approach of juxtaposing the effects of sleep on four gists, each reflecting an apparently different gist extraction operation: inferential gist, which requires extraction of relations across non-contiguous events; statistical learning, in which regularities must be extracted from a series; summary gist (which includes the previously mentioned false memory), in which a theme is abstracted from a temporally contiguous series; and category gist, requiring characterization of a stimulus at a higher level in the semantic hierarchy. We define these four gists in Box 1, and provide a conceptual depiction of the tasks we used to measure them in Figure 1. A visual depiction of trials in each task is given in Figure 2. Although the summary gist that we chose in this task is false memory, other forms of summary gist may include the semantic summary of a group of pictures or a series of objects located in a single picture. We do not claim that these four are the only “gists”, but rather that these are a subset we selected for this study. We selected these in particular, because they differed across a number of different parameters, including temporal continuity and gist extraction of a pattern or a single instance. In this study, we could only cover one type of summary gist (commonly referred to as false memory), but we predict that other forms of summary gist would act in a similar manner, based on the common property of reflecting a high-level summary. We reasoned that the role of REM or SWS in gist memory would be best elucidated by comparing the effect of these sleep stages on different gists. We anticipated that each gist would either increase over time, or decay at a slower rate than our reference measures, detail memory.

**Figure 1.**
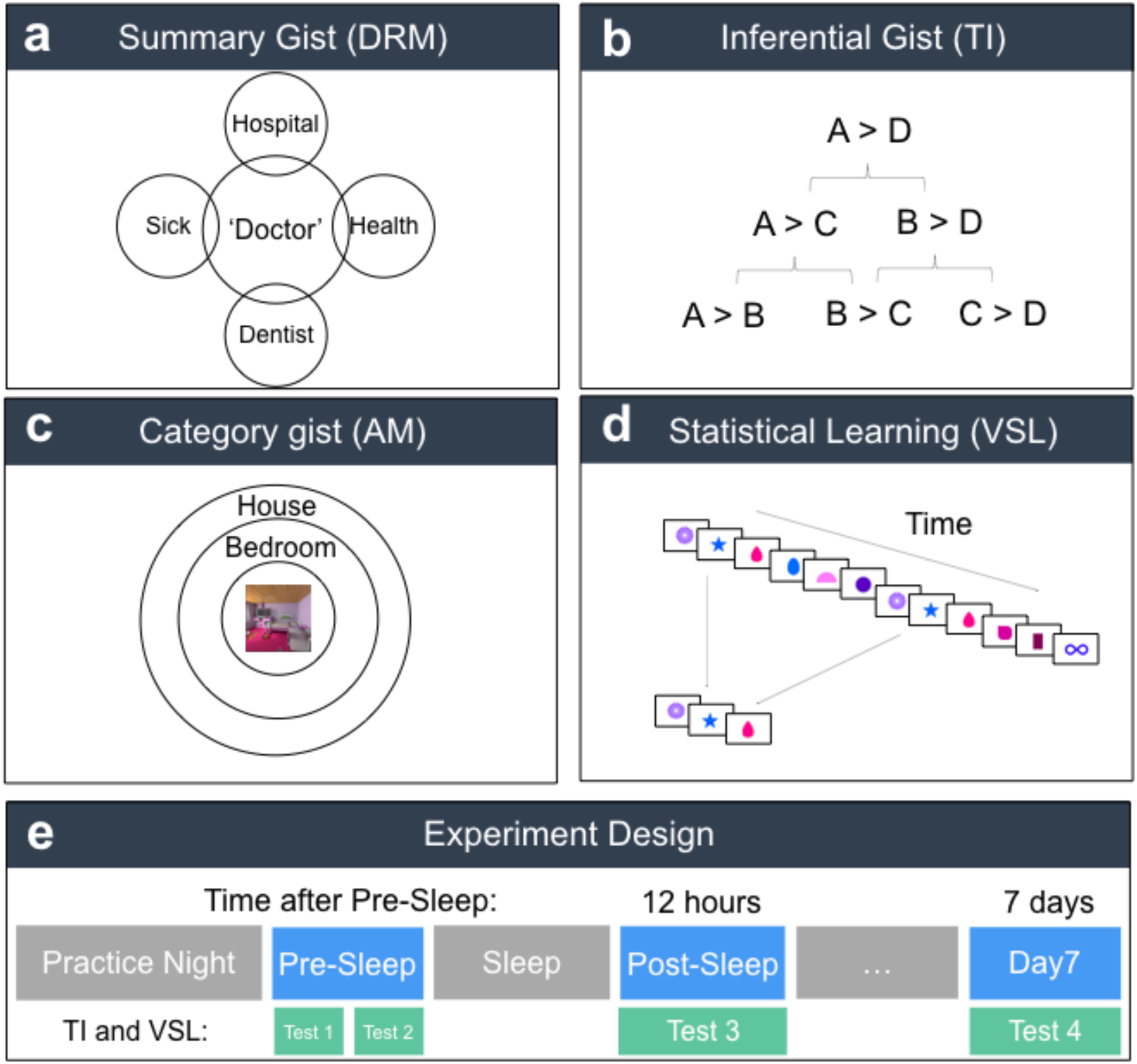
Multiple conceptualizations of gist. a) Conceptual illustration of overlapping meaning between studied words and the unstudied “gist” word in the DRM task. b) Conceptual illustration of studied premise pairs on the bottom row, unstudied first-order inferences on the second row, and unstudied second order inferences on the top row. c) Conceptual illustration of gist internal to an individual stimulus. Participants studied word-scene pairs, and were then asked about the scene’s category (e.g., bedroom) and super-category (e.g., house). d) The left-hand panel shows a sequence presented over time, and one triplet which is repeated in the sequence. e) Study design. Participants first completed a practice night with the Sleep Profiler. Then they completed a study and first test session (Pre-Sleep test), which included two test-retests, Test 1 and Test 2 for TI and VSL tasks. Twelve hours and 7 days later participants completed a second (Post-Sleep Test) and third (Day7) test session, respectively.

**Figure 2.**
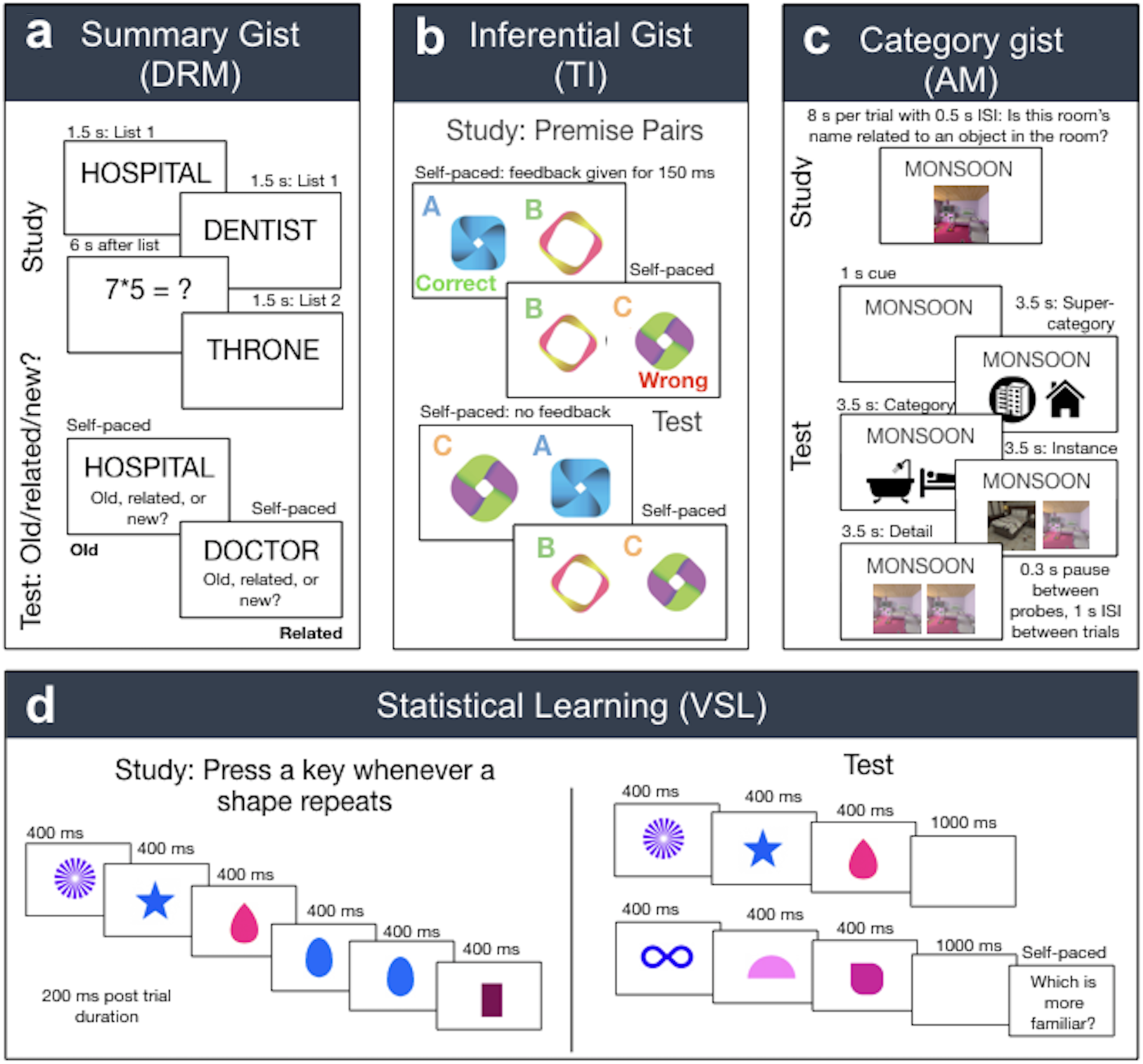
Trials in each task during study and test. a) In TI, participants were presented with self-paced study trials. After a shape was selected, feedback was displayed at the bottom of the screen for 150 ms. During test, trials were also self-paced with no feedback provided. b) In the DRM task, participants were presented with a list of words at 1.5 s per word. After each list, participants solved math problems for 6 seconds. During test, participants indicated whether each word was old, related, or new in self-paced trials. c) In the AM task, participants completed incidental encoding trials, they were asked to study each word-scene associate for 8 seconds to determine whether the room’s name could be plausibly related to an object in that room. We selected this format to encourage participants to look at all aspects of the room, including details. During test, participants were cued with the word associate for 1 second, and then answered a cascade of forced-choice probes for each associate at four levels: super-category, category, instance, and detail. Each probe was presented for 3.5 seconds, with a 0.3 second pause between probes and a 1 s inter-stimulus interval between trials. d) In the VSL task, participants were shown a stream of shapes at 400 ms intervals with 200 ms post trial. They were asked to press a key whenever a shape repeats. During test, participants were shown two triplets of shapes at 400 ms per shape and 1s between sequences, and then asked which was more familiar.

We also took an individual differences approach: because false memory, transitive inference, and statistical learning have all been shown to have sleep effects compared to wake, we wanted to probe these effects deeper by looking only at individual differences in sleep stages. The IOtA account^11^ and associative linking during REM theory^8^ provided theoretical frameworks for gist extraction in general that could implicate any of our tasks. Therefore, we predicted that participants with either more SWS or REM sleep would have better gist memory. Due to a separate line of research proposing that the anterior hippocampus is specialized for gist memory and the posterior for detail,^14^ we also collected structural MRI data. We predicted that larger anterior hippocampal volumes would be associated with better gist memory. We pre-registered our hypotheses on the Open Science Framework (OSF^15^; see osf.io/kqc7z). Although some of our hypotheses were open-ended in that multiple gist measures were related to either SWS or REM, our pre-registration did significantly constrain our analysis in preprocessing, included variables, and decision rules, among others.

## Materials and Methods

### Participants

104 participants were recruited in Kingston, Canada using posters, Facebook advertisements and web posts on Reddit, Kijiji and Craigslist. Participants were required to be: between 22-35 years of age (to avoid potential developmental effects); right-handed, an English native-speaker, have normal or corrected-to-normal vision and hearing, typically sleep at least 5 hours a night, have no contraindications for MRI scanning, have no history of neurological disorders, sleep disorders, or recurrent mental illness that included medication, and not currently be taking psychotropic medications.

Recruits meeting these criteria were invited to undergo a two hour in-person screening session in which demographic eligibility was confirmed, and a simulated MRI scanner was used to rule out claustrophobia, inability to keep still (operationalized as falling below the 95^th^ percentile of low-frequency motion as measured in a reference sample), and inability to stay awake for a 20-minute eyes-open resting session. In addition, we assessed participants’ ability to respond to at least 33% of encoding trials in a recognition memory task, to obtain a d-prime of at least 0.1 in a subsequent memory test, and to demonstrate acceptable reading comprehension and speed (a TOWRE score of at least 26.4 and a Nelson-Denny score of 2). Finally, participants were excluded for multiple no-shows, failing to follow instructions, rudeness, and excessive drug use. Participants were advised that they would be compensated CAD$20 for their time (or a pro-rated amount in the case of early withdrawal).

Of the original sample, 69 participants met our criteria and returned to complete the full experiment. The average age of participants who did so was 26.83 years (*SD* = 4.21 years). They described themselves as men (*n* = 30), women (*n* = 38), and other (*n* = 1). We used G*Power 3.1 to calculate a sensitivity power analysis, and found that based on our sample size of 69 participants and models with 3 predictors, we had 80% power to detect a small (0.09) effect size.

### Experimental Design and Statistical Analysis

After their screening session, participants wore a sleep EEG device (Sleep Profiler, Advanced Brain Monitoring, Carlsbad, CA; see Fig. 3A) for a habituation night, during which they became accustomed to wearing and operating it. After the habituation night, on the basis of the recorded data, participants received corrective training as necessary.

**Figure 3.**
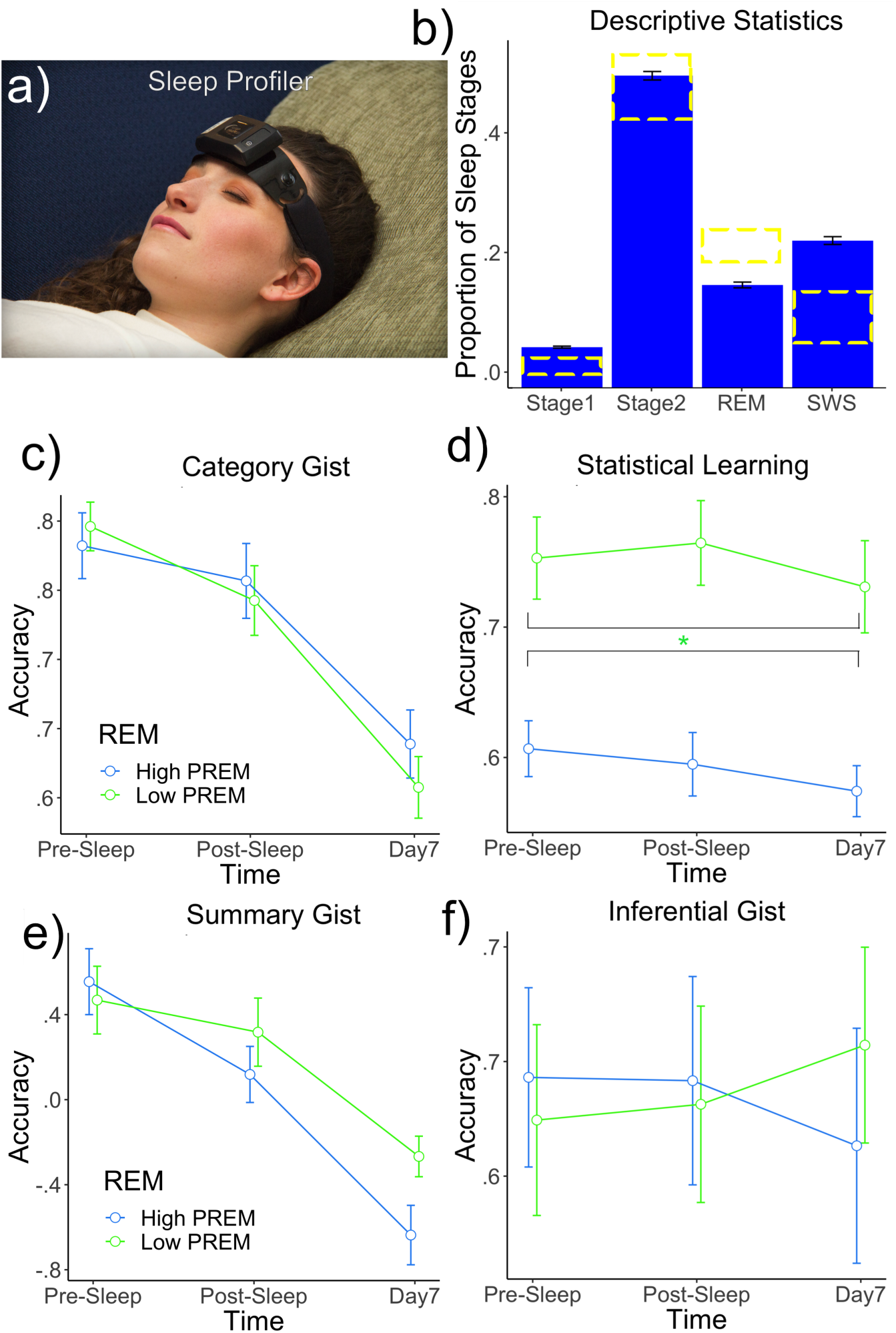
Proportions of different sleep stages and proportion of REM effects on memory. a) Photograph of the Sleep Profiler, a single-channel EEG headband with dual EOG that participants took home to record their sleep. b) Barplot of proportions of different sleep stages in this dataset. Error bars reflect standard error. To assess whether populations of sleep stages in this dataset were close to population norms, we included yellow boxes that indicated proportions of sleep stages among healthy young adults as described by Carskadon and Dement^54^. c-e) For visualization purposes, we median-split REM into two groups, though our analysis was continuous. High and low REM groups across statistical learning, associative gist, and inferential gist measures. Proportion of REM significantly negatively predicted statistical learning collapsing over or controlling for time.

Participants conducted an initial study phase during an evening visit, which was followed by a test phase (Pre-Sleep test). After sleeping at home while measuring their own sleep using a take-home sleep EEG device, they returned the next morning (12 hours after their initial session) for another memory test (Post-Sleep test). A week later, they returned for a final memory test (Day7). They also completed an MRI session several weeks after testing.

The study session and each test visit included a study or test phase from each of four memory tasks (Fig. 2). These included a transitive inference (TI) task (inferential gist), a Deese, Roediger and McDermott (DRM) task (summary gist), a visual statistical learning (VSL) task (statistical learning), and a word-scene associative memory task (category gist). These tasks are described in detail below. After each study and test session, participants were encouraged to take a break. In addition, we administered an operation span task (OSPAN) at each session to measure participant fatigue.

As the data collection effort involved collaboration with researchers investigating a variety of individual differences within our participant group, our participants also completed further experimental sessions involving objectives unrelated to the current study goals (to be reported elsewhere).

As pre-registered, we excluded data falling three median absolute deviations above or below the mean of each variable. To investigate relationships among gist and detail variables, we ran Pearson correlations. To investigate predictors of patterns of change in gist and detail over the course of a week, we used multi-level modelling (MLM) implemented in the nlme package in R.^16, 17^ Multi-level modelling allowed us to estimate an individual change function in each participant, as well as predictors of this change function. Therefore, this analysis is especially suitable for looking at individual differences in change over time.^18^ In the models, we estimated each participant’s change over three time points at Level 1 and individual differences at Level 2. Our predictors were Time, as well as Level 2 predictors related to individual differences in sleep and hippocampal volumes. We first ran random intercept and random slope models, and if this model did not converge, we ran random intercept only models. Time was coded as 0 for Pre-Sleep, 1 for Post-Sleep, and 2 for Day7. In all multi-level models, we included grand-mean centered hippocampal volumes (aHPC and pHPC) or sleep stages (proportion of SWS or REM) as predictors. We also included fatigue (OSPAN) as a control variable. Outcome variables were measures of gist memory on four different tasks at 3 time points: in the same evening as study (Pre-Sleep), the next morning (Post-Sleep), and one week later (Day7). To balance control for multiple comparisons with experimental power, we selected a 0.05 False Discovery Rate for each independent variable.^19^ Lastly, to investigate whether gist and detail measures decayed at different rates, we conducted two MANOVAs. For the AM and DRM tasks, we conducted a MANOVA for 3 time points and 3 dependent variables, followed up by ANOVAs. For the TI task, we conducted a MANOVA for 3 time points and 2 dependent variables, followed up by ANOVAs.

### Test-Retests and Interference

We took a number of steps to mitigate possible inference across and within our tasks. We selected different types of stimuli in different tasks (simple shapes in VSL, scenes in the AM task, words in DRM, and complex shapes in TI) to reduce cross-task interference. In TI and VSL tasks, participants needed to learn complex relationships among items to be tested at multiple time points. We could not test a different set at each time point, as the large number of relationships involved could generate high interference and require excessive study time. Instead, for VSL we used one set by Turk-Browne et al. ^20^ (which we called compound shapes) and generated a distinctive set (which we called simple shapes, see osf.io/kqc7z/ in VSL stimuli folder) in a way designed to assess possible test-retest effects (see Fig. 1). The compound shapes stimuli were provided by the *Millisecond Test Library*,^21^ but we changed the colours from red and green to different colours for each shape. For TI we used two sets of symbols from collections of abstract vector symbols online (which we called circles and transparent shapes). Section A (randomly assigned to either compound shapes or simple shapes for each participant for VSL and circles or transparent shapes for TI) was tested twice at the Pre-Sleep test, once at the Post-Sleep test, and once at Day7. Section B (randomly assigned to either compound shapes or simple shapes for VSL and circles or transparent shapes for TI) was tested once at each test session. Due to an error, for three participants, more than 7 sessions were administered. In these cases, we measured only the first administration.

In the TI task, there were no test-retest effects for premise pairs, first-order inferences, or second-order inferences between the two tests at the Pre-Sleep session, *p*s > .407, suggesting any longitudinal trends were unlikely to be attributable to repeated testing. Also, both sets of stimuli (circles and transparent shapes) were independently above chance for all time points, *p*s < .001. In the VSL task, simple shapes were also above chance at all time points, *p*s < .001, but compound shapes were not at the Post-Sleep test, *p* = .55, or Day7 test, *p =* .58. The statistical learning task was divided into determinstic and non-deterministic sequences, which we will describe in detail in the following section. There was a significant test-retest effect in memory for deterministic sequences from simple shapes, *t*(44) = −-2.09, *p* = .043, *r* = .86, indicating that as participants were tested on the material a second time during Pre-Sleep test, their scores improved. There was no significant test-retest effect in memory for deterministic sequences from compound shapes, *t*(24) = 0.883, *p* = .386, *r* = .657. There was no significant test-retest effect in memory for non-deterministic sequences from simple shapes, *t*(44) = 0.06, *p* = .951, *r* = .72 or compound shapes, *t*(24) = 1.528, *p* = .140, *r* = . 742.

### Tasks

#### Visual statistical learning (VSL)

VSL^20^ provides a measure of statistical learning, which has been defined as implicit learning of patterns that are automatically segmented from a continuous environment (such as co-occurring shapes in a sequence^22^). To implement this task, we modified a publicly-shared script (described by Turk-Browne et al. ^22^) obtained from the *Millisecond Test Library*.^21^ During learning, participants were shown four blocks of a stream of shapes over 312 trials (each appearing for 400 ms with 200 ms post-trial duration) and instructed to press a key whenever a shape repeated. Two blocks were shown for each stimulus set, which were interweaved. At test, in each of 64 trials, participants were presented with two short series of three shapes familiar from the learning phase (each appearing for 400 ms with an inter-series interval of 1s) and instructed to select the temporal sequence that seemed more familiar. Some triplets were sequences seen in the previous learning task (e.g., ABC), and others were novel sequences (e.g., AEG). In our adaptation of this task, half of target triplets involved shapes that always appeared in the same order during training (i.e., deterministic sequences; e.g., ABC), whereas the rival sequence was never encountered (e.g., BAC). The other half of triplets sometimes also appeared in the rival sequence (i.e., non-deterministic sequences; e.g., ABC was presented at a 3:1 ratio to BAC). The sequence that was presented more frequently was always considered the correct answer.

#### Transitive inference (TI)

The TI task^4^ provides a measure of inferential gist, or the ability to make inferences across paired associates that form an implicit hierarchy (e.g., A > B > C > D). To implement this task, we again modified a script (described by Frank et al.^23^) obtained from the *Millisecond Test Library*.^24^ During study, participants were introduced to the hierarchy by being presented with four pairs of symbols (i.e., “premise pairs”; e.g., A and B, B and C, C and D, D and E), and asked to guess the winning symbol (e.g., A > B, B>C, C>D, D>E; Fig. 1B and Fig. 2B). For each premise pair, the winning symbol was counterbalanced on the left and right sides. Participants were instructed to first find the winning symbol by trial and error, and then over a number of trials they would be able to learn the correct winning symbol with the provided feedback. Participants were given feedback for 150ms about the correctness of their guess after each study trial. If the participants selected the correct winning pair, green text that said ‘correct’ would appear at the bottom of the screen. If they selected an incorrect pair, red text that said ‘incorrect’ would appear. Participants indicated their responses by pressing ‘A’ for left-side stimulus and ‘L’ for right-side stimulus on a standard keyboard. Items were organized into blocks, each block containing 100 trials. Each block presented each of 5 pairs of symbols in 20 trials (10 trials forward and 10 trials backward) randomly. Items within each block were arranged in pseudo random order to avoid revealing the hierarchy. The same premise pair was never presented twice in a row, and the neighbouring pairs never followed one another (e.g., AB, BC, CD, DE, EF). Participants were trained to a criterion of 0.75 that was dynamically evaluated throughout training and calculated at the end of each block. An additional block of 20 trials (4 for each pair) was presented in the participants did not reach criterion to a maximum of 5 additional blocks.

Participants studied two stimulus sets, which were divided by a mandatory 12 second break with an option to resume anytime thereafter. At test, participants completed 180 test trials without feedback testing both the original premise pairs as well as inference pairs (e.g., B > D). Participants completed 20 test trials per premise and inference pair: these included five premise pairs, two first order inference pairs (B > D and C > E), one second order inference pair (B > E), and one non-inference pair (A>F). Test trials advanced following a response by the participant. Inference pairs are higher-order relationships that must be inferred from the relationships between lower-order relationships (e.g., correctly identifying that A > C would require inference from the premise pairs A > B and B > C). Correctly inferring such higher order relationships is taken as evidence for the formation of a superordinate hierarchy.

#### Word-scene associative memory (AM)

The goal of the AM task was to provide gist and detail memory measures that were internal to an individual stimulus, rather than linked across a series or set of items. Our approach was to use single words to cue scene associates (i.e., word-scene pairs) that could be characterized in various levels of detail. For the encoding task, participants viewed each word-scene pair for 8 s under incidental learning instructions designed to bind the word cue and picture associate: in particular, to decide whether the “name” each room had been given could plausibly have been assigned based on an object found in that room. A 0.5 s ISI separated each trial. After the encoding task was complete, we probed four measures of memory for each studied item: super-category (general semantic category of scene, e.g., “hotel”), category (room type within category, e.g., “lobby”), instance (specific exemplar of a lobby), and detail (exact contents of the scene). During test, participants were cued with the word associate for 1 s, then answered a cascade of two-option forced-choice questions about its associate, deciding on whether it is a word associated with a noise or room image, super-category, category, instance, then details. Super-category and category levels for each probe were presented both verbally and visually: e.g., house, restaurant and school icons. Instance was tested by presenting two scenes from the same category and details were tested by presenting two arrangements of the same scene with different details. In the instance probe, all options were shown using a variant of the images that excluded the feature to be assessed in the final “detail” probe, so as not to reveal the correct answer in that final stage. Importantly, there was only one encoding trial, so higher-level categorical representations would have to be inferred from the same representation that contains the instance itself. Each probe in this series of four probes was presented for 3.5 s with a pause of 0.3 s between each probe, and a 1 s ISI between each trial. For each multiple-choice question, even if participants responded to the previous level incorrectly, lures were selected within the correct answer to the prior probe (e.g., if the correct super-category of an associate to a word was “house”, the options for the category probe would be *bathroom, bedroom* and *kitchen*). This was in order to account for possible independence between these memory types, for instance participants might remember a detail arrangement but not the scene’s category.

Scene associates belonged to three super-categories (e.g., *house, restaurant, school*), which were all divided into 3 categories (e.g., *bathroom, bedroom and kitchen* for the ‘house’ category). Each category contained 3 instances (e.g., the room-type category *kitchen* contains 3 separate kitchens). Finally, each instance contained 3 detail variants. One detail image was the one originally studied and the other two have been edited to replace one object with a novel object. During each of the three test sessions, we tested participants on three previously-untested super-categories to avoid test-retest effects.

#### Deese, Roediger and McDermott (DRM)

The DRM task provides a measure of summary gist, the ability to extract meaning common to a set of related items. The difference between summary gist and inference gist is described by Stickgold and Walker^25^ as being akin to the difference between sets and relations, respectively. To implement this task, we again modified a publicly-shared script obtained from the *Millisecond Test Library*.^26^ During study, participants were asked to learn 27 lists of 14 related words each. Words were presented with a stimulus onset asynchrony of 1.5 seconds, with each list separated by 6 s break during which participants were asked to solve simple math problems. During test, participants were presented with 27 target words (old words from the study list), 9 critical lures (related words associated with the theme of each word list) and 36 unrelated lures (new words from an unstudied category). These words were presented one at a time in random order and trials only progressed when participants made a response, selecting among ‘remember’, ‘related’, or ‘new’.

In analysis of this task, our goal was to derive gist and detail measures from the 3×3 matrix of old, related and new responses to old, related, and new items. Stahl and Klauer used response rates to model group estimates of gist and detail^27^; however, this approach failed when running models for individual participants. We instead modelled summary gist using a d’-based analysis, drawing inspiration from Dandolo and Schwabe,^28^ who described two relevant models. An “Old Distinct” model expects old memory representations to be distinct from both related and new ones, indicating a precise detailed memory; and an “Old and Related Similar” model expects old and related memory representations to be quite similar, while being distinct from new items (suggesting that gist has been encoded but detail has not). Because one model captures only detail, and the other only gist, the two are orthogonal. We adapted this framework to our task: in the “Old Distinct” model, hits were the proportion of old responses to old items, and false alarms were the proportion related responses to old items (we refer to this model as word detail). In the “Old and Related Similar” model, hits were the proportion of old or related responses to old or related items, and false alarms were the proportion of new responses to old or related items (we refer to this model as summary gist). To assess the latter model, we collapsed old and related rows and columns from our 3×3 matrix, yielding a 2×2 matrix of old+related and new responses to old+related and new items for each participant. Finally, we replaced all values of 0 and 1 in the resulting 2×2 matrix with 0.00001 and 0.9999 in order to avoid hit or false alarm rates of 0 or 100%. Then, we calculated d’ values for both the “Old Distinct” (memory for individual words) and “Old and Related Similar” (summary gist) models.

#### Operation span (OSPAN)

Lopez et al. found that the simulated flight performance of sleep-deprived Air Force pilots was predicted by OSPAN.^29^ Hence, it is often used as a suitable proxy for performance-related fatigue. Here, we used the OSPAN task to measure and control for variability in participant alertness across each test phase, using a task described by Unsworth et al.^30^ and obtained from the *Millisecond Test Library*.^31^ The task consisted of 5 practice letter questions, 15 practice math questions and 15 testing trials. Practice trials had no time limit and featured a 0.5s ITI. For practice letter questions, three to seven letters were shown in series on the screen. Each letter was presented for 1 second with a 200ms ISI. After presentation, participants were asked to enter the letters that they had seen in the correct sequence on the onscreen keyboard. Participants received feedback on their answers for 2 seconds. For practice math questions, participants were asked to complete math operations (e.g., (1*2)+1). They were instructed to complete the math problem as quickly as possible. Once they had an answer, they clicked the button to progress to the next screen. A possible answer was displayed on screen and participants were asked to indicate whether the answer is ‘true’ or ‘false’. Practice trials were used to calculate the mean time that it takes a participant to solve math problems. For test trials, letter and math questions were intermixed, and there was a limit of the participant’s average math problem time plus 2.5 SDs before the trial progressed on its own (to discourage letter rehearsal). Each test block consisted of both letter and number trials. There were three blocks of each of the five set sizes (i.e., there were three repetitions of 3, 4, 5, 6, and 7 letter and number sizes). The shortened OSPAN consisted of five practice letter questions, five practice math questions and one block of each of four set sizes (i.e., one repetition of 3, 4, 5, and 6 letter and number sizes).

### Apparatus

Participants completed three sessions individually in a testing room. One task was executed using MATLAB with Psychtoolbox^32^ and SuperPsychToolbox.^33^ The remaining three tasks (OSPAN, DRM, VSL, and TI) were executed using Inquisit (Version 5.0.6.0^34^). For sleep stage measures, we used the Sleep Profiler, a single-channel electroencephalography (EEG) device worn on the forehead that records at 256 Hz from three sensors at approximately AF7, AF8, and Fpz^35^ (Sleep Profiler Scoring Manual, 2015). We chose to use the Sleep Profiler over a full polysomnography net (PSG) because it enabled us to measure more naturalistic sleep, since it is designed to be used independently by participants at home, whereas PSG measurement typically requires that participants sleep in a sleep lab. Use of Sleep Profiler devices also allowed us to collect more data, since they do not require time-intensive monitoring or manual sleep-staging. The device applies a 0.1 Hz low-frequency filer and a 67 Hz high-frequency filter. Expert review of Sleep Profiler data and concurrently-collected data from a PSG (which is broadly regarded as a “gold standard” for sleep data collection) has previously resulted in strong agreement for total sleep time (*ICC* = 0.96) and REM sleep (*ICC* = 0.92), and poorer agreement for Stage 1 (*ICC* = 0.66) and Stage 3 (*ICC =* 0.67).^28^ Agreement for Stage 3 increased when combined with Stage 2 into a non-REM (NREM) measure (*ICC* = 0.96).

The Sleep Profiler system also features automated sleep-staging software, which we manually validated using a subset of raw sleep data (described below) gathered using our Sleep Profiler. Briefly, as described by Levendowski et al.,^36^ the software first rejected 30-second epochs where the absolute amplitude was ≥ 500 µV, applied a notch filter, and then an infinite impulse response band pass-filter to obtain 16 Hz samples of the power values for delta (1-3.5 Hz), DeltaC (delta power corrected for ocular activity), theta (4–6.5 Hz), alpha (8–12 Hz), sigma (12–16 Hz), beta (18–28 Hz), and EMG bands (> 40 Hz with a 80 Hz, 3 dB rolloff). Another set of power values was derived after application of a 0.75-Hz high-pass filter. Both filtered and unfiltered power spectra were used to stage sleep. If at least 15 s of valid data was available, AF7-AF8 channels were used for scoring, followed by AF7-Fpz and AF8-Fpz. When either of the latter were used, power spectra were increased to compensate for signal attenuation due to shorter interelectrode distances.

Power spectra averaged from 16 to 4 Hz were used to detect sleep spindles, which were characterized by spikes in absolute and relative alpha and sigma power that met empirical thresholds. Spindles were at a minimum 0.25 Hz in length with no maximum. To reduce misclassifying spindles, beta and EMG power bands had to be simultaneously surpassed relative to the alpha and sigma power.

### MRI data acquisition

The day prior to each participants’ MRI scan, they completed a biofeedback session in the simulated MRI scanner to become better habituated to an MRI-like environment and learn to reduce their head movement (and thereby improve the signal quality of their brain images sampled the next day). During the biofeedback session, participants viewed a 45-minute documentary with a live readout of their head motion overlaid. When their head motion exceeded an adaptive threshold, the documentary was paused for several seconds while static was played on the screen along with a loud, unpleasant noise. Following the documentary, a brief memory test was administered outside of the mock scanner to ensure participants were paying attention to the film content rather than just their motion (not analyzed here).

The next day, we used a whole-body MRI scanner (Magnetom Tim Trio; Siemens Healthcare) to gather a variety of image sequences over the course of a 1.5 hour scan. As described in our pre-registration, our hypotheses in the current study related specifically to the medial-temporal lobe anatomical analyses. To assess these predictions, we gathered high-resolution whole-brain T1-weighted (T1w) and T2-weighted (T2w) anatomical images (in-plane resolution 0.7 x 0.7 mm^2^; 320 x 320 matrix; slice thickness 0.7 mm; 256 AC-PC transverse slices; anterior-to-posterior encoding; 2 x acceleration factor; *T1w* TR 2400 ms; TE 2.13 ms; flip angle 8°; echo spacing 6.5 ms; *T2w* TR 3200 ms; TE 567 ms; variable flip angle; echo spacing 3.74 ms) and an ultra-high resolution T2-weighted volume centred on the medial temporal lobes (resolution 0.5 x 0.5 mm^2^; 384 x 384 matrix; slice thickness 0.5 mm; 104 transverse slices acquired parallel to the hippocampus long axis; anterior-to-posterior encoding; 2 x acceleration factor; TR 3200 ms; TE 351 ms; variable flip angle; echo spacing 5.12 ms). The whole brain protocols were selected on the basis of protocol optimizations designed by Sortiropoulos et al..^37^ The hippocampal protocols were modelled after Chadwick et al..^38^

### Hippocampal volumes

As noted in our pre-registration, we did not have any specific hypotheses regarding laterality for hippocampal variables. Therefore, before beginning data analysis, we decided to perform a Pearson’s correlation of aHPC and pHPC volumes across hemispheres. If the correlations were greater than 0.9, we would merge the volumes across hemispheres. Otherwise, we would run them separately in our models. aHPC volumes across hemispheres had a correlation of 0.686, and pHPC volumes across hemispheres had a correlation of 0.458. Therefore, we analyzed left and right hippocampal volumes separately in all of our models, investigating only either left or right hippocampal volumes in any given model.

The ultra-high-resolution T2w 0.5mm isotropic medial temporal lobe scans were submitted to automated segmentation using HIPS, an algorithm previously validated to human raters specialized in segmenting detailed neuroanatomical scans of the hippocampus.^39^ Three independent raters (including one of the authors) were trained on segmenting the hippocampus at the uncal apex into aHPC and pHPC segments, and achieved a Dice coefficient of absolute agreement of 80%. Two of these raters (including one of the authors) independently segmented all participants in this study using the 0.5mm T1w scans. The T2w medial temporal lobe scans were registered to the T2w whole-brain scans, which were in turn registered to the T1w whole-brain scans, and the combined transform was used to place the rater landmarks on the detailed medial temporal lobe scans. All voxels belonging to the hippocampus posterior to this coronal plane containing this point were classified as posterior hippocampus (pHPC), whereas all points anterior to and including this plane were classified as anterior hippocampus (aHPC). Finally, the total number of voxels in each subregion was multiplied by the volume of each voxel to obtain a total aHPC and pHPC volume. These steps were followed separately in each hemisphere. Freesurfer segmentation of the T1w and T2w brain data (v6.0^40^) was used to compute Estimated Intracranial Volumes, which in turn were used to control for effects of overall head size in the hippocampus volume vectors.

### Sleep Profiler validation

We selected a random subset of ten participants and used their raw sleep EEG data for validation of the Sleep Profiler auto-staging. Two independent raters rated the data and we took the average of their ratings. Prior to scoring, we pre-registered on OSF that if intra-class correlations (ICCs) were good (between 0.75 and 0.9) or excellent (over 0.9) for SWS, REM sleep, and TST^41^, we would use the sleep stage values derived from the automated scoring system. If they were below 0.75, then we would manually score the sleep data. We used an ICC measure that estimates the degree of consistency across measurements rather than absolute values.^42^

ICCs between the automated scoring system and two independent raters, as well as inter-rater ICCs, are given in Table 1. Human rater agreement with one another was excellent for TST and REM, whereas their agreement with the algorithm was lower, at a level closer to 0.8 (good). Human rater agreement with the algorithm was also good for SWS, whereas SWS agreement among human raters was only moderate. Having observed consistently good algorithm classification performance (with all ICCs above a cutoff of 0.75), we therefore used the sleep stage values derived from the automated scoring system for all of our analyses. We did, however, perform a manual quality control inspection of the sleep data to remove 1) any recordings that contain no SWS or no REM, 2) partial recordings that did not contain the full night of sleep, and 3) recordings where a participant’s total sleep time was less than 50% of how much they typically reported sleeping on weekdays. This resulted in the omission of data from four participants.

**Table 1.** Intra-class Correlations between the Sleep Profiler Automated Scoring System and two Independent Raters.

| Sleep stages | Intra-class correlations (ICCs) | Inter-rater ICCs |
| --- | --- | --- |
| TST | 0.855 | 0.982 |
| SWS | 0.819 | 0.669 |
| REM | 0.798 | 0.988 |

TST = Total Sleep Time, SWS = Slow-wave Sleep, REM = Rapid Eye Movement Sleep.

## RESULTS

The goal of our study was to investigate the influence of sleep, including specific sleep stages, on different forms of gist, juxtaposing those requiring rule extrapolation from relations against those requiring gist extraction from sets. We approached this question both within-subjects – evaluating relative change in memory over time – as well as between-subjects, evaluating the predictive power of sleep variables over gist extraction. We began our between-subjects analysis by investigating correlations among our memory variables to identify possible correspondence among memory types. We then compared gist and detail memory variables within the same tasks using a MANOVA. Afterwards, we investigated patterns of change in gist and detail memory as a function of time and sleep stages. As pre-registered, we also ran a version of the model incorporating hippocampal volumetric predictors, but as these were not predictive of memory performance, we re-ran our model without a hippocampal factor and present it in this form for simplicity. We investigated models with both interactions over time, as well as models controlling for time. Descriptive sleep statistics are given in Figure 3B. Before addressing our research questions, we first discuss baseline models.

### Memory predictors

We analyzed only variables in which participants achieved above-chance and off-ceiling performance as of Pre-Sleep, to ensure that our measures were sensitive to change in both a positive and negative direction. Memory accuracy for all variables at Pre-Sleep is given in Table 2. One-sample t tests showed that all of the outcome variables in AM, TI, and VSL were significantly above chance and below ceiling as of Pre-Sleep, all *p*s < .001. In the DRM task, one-sample t tests showed that memory for summary gist was above chance for Pre-Sleep, *t*(69) = 2.960, *p* = .004. However, at Post-Sleep and Day7, summary gist was not significantly above chance. Word detail memory was not above chance for any time points, so we excluded this variable from subsequent analyses.

**Table 2.** Memory performance at Pre-Sleep.

| Memory Variable | Mean accuracy | Standard Deviation |
| --- | --- | --- |
| Premise Pairs | 0.698 | 0.155 |
| First Order Inferences | 0.603 | 0.201 |
| Second Order Inferences | 0.635 | 0.258 |
| Deterministic Sequences | 0.623 | 0.154 |
| Non-Deterministic Sequences | 0.618 | 0.141 |
| Super-Category | 0.774 | 0.129 |
| Category | 0.817 | 0.127 |
| Instance | 0.915 | 0.094 |
| Detail | 0.665 | 0.122 |
| Summary Gist | 0.351 | 0.906 |

To explore possible relationships and redundancies among our behavioural memory measures, we inspected a correlation matrix, focusing on relationships involving measures from the same task (Fig. 4A), and to a more limited extent, across tasks. Because the goal of this step was to limit our predictors rather than form inferences about tasks, we did not apply multiple-corrections comparisons in this step. As there were strong correlations in all tasks, to simplify our analyses, we combined gist measures within our tasks: first-order and second-order inference memory into an inferential gist measure; non-deterministic and deterministic sequence memory into a statistical learning measure; super-category and category into a category gist measure. As a result of this step, the remainder of our analyses concerned only four variables: summary gist, category gist, inferential gist, and statistical learning. Memory performance for all gist variables at Pre-Sleep is given in Figure 4B.

**Figure 4.**
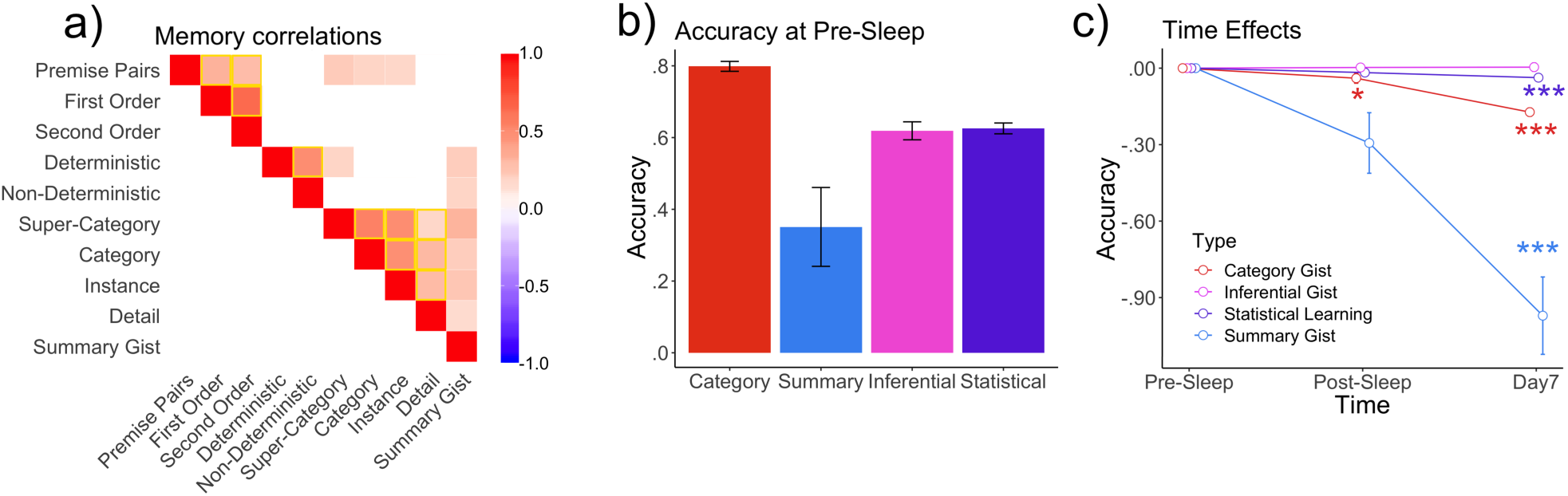
Memory variables at Pre-Sleep, Post-Sleep, and Day7. a) Correlations among dependent variables averaged over time. Yellow boxes indicate potential relationships among variables that are within the same task. Missing values indicate non-significant relationships. b) Accuracy at Pre-Sleep for all types of gist. c) Time effects for all four dependent measures. All variables were normalized to 0 at Pre-Sleep only for visualization purposes, and not in the statistical analyses. Error bars indicate standard error. Some error bars are too small to be visible. Category gist significantly decreased from Pre-Sleep test to both Post-Sleep test and Day7. Inferential gist was retained from Pre-Sleep test to both Post-Sleep test and Day7. Statistical learning and summary gist significantly decreased from Pre-Sleep test to Day7. Only significant p values after FDR correction are indicated. *** indicates *ps* < .001, * indicates *ps* < .05.

Special consideration is owed to collapsing of the VSL task. Fuzzy Trace Theory, one of the prominent frameworks distinguishing between gist and detail memory, defines detail as information that was tested in the exact form it was learned.^1^ Given this definition, we initially felt deterministic sequence memory could be considered a detail measure and non-deterministic sequence memory a gist measure, which is the framework we followed in our pre-registration. However, as both deterministic and non-deterministic sequences require extracting patterns from a sequence, and as we found them both to be correlated in the analysis above, we feel they are both better-represented as gist measures. Non-deterministic sequences additionally require participants to sift through more noise than deterministic sequences, so can be considered higher on a hierarchy of gist. A better detail measure for the task would have been a measure of participants’ memory for individual shapes (not measured in the current analysis).

Hence, we were left with four putative detail memory predictors: “instance” and “detail” probes in our AM test (scene-cued object recognition), memory for premise pairs (TI recognition of explicitly studied shape relations), and individual word recall (DRM). As we excluded individual word recall from analysis due to low performance (as discussed above), We were left with the AM and TI measures only. We performed MANOVA analyses to assess divergence of gist and detail over time for these two tasks. Category gist decreased by 22%, whereas instance memory (AM) decreased by 15%, and detail (AM) by 6%. For AM, there was a significant MANOVA for Time differences across category gist, instance and detail memory, *F*(2,182) = 13.176, *p < .*001, η2 = 0.179. Follow-up ANOVAs indicated significant decay rates in category gist, *F*(2,182) *=* 38.721, *p* < .001, and instance memory, *F*(2, 182) = 28.496, *p* < .001, but not detail memory, although a trend was revealed in this direction, *F*(2, 182) = 2.804, *p* = .063. Inferential gist increased by 0.6%, whereas premise pairs decreased by 2%. For TI, the MANOVA for time differences across premise pair memory and inferential gist memory was not significant, *F*(2, 205) = 0.219, *p* = .928, η2 = 0.002.

### Baseline Multi-Level Models

We next constructed a baseline model to compute Intraclass Correlation Coefficients, which indicate the amount of variance in dependent variables that can be attributed to individual differences (and that can therefore be reasonably attributed to individual difference predictors, such as sleep). Variables with 10% or more individual differences variance are considered to have meaningful individual differences. The ICCs were *ρ* = 0.848 for statistical learning (VSL), *ρ* = 0.085 for category gist (AM), *ρ* = 0.110 for summary gist (DRM), and *ρ* = 0.875 for inferential gist (TI). This means that, across different memory measures, between 9 and 88% of the variance in memory variables can be attributed to individual differences, although the low intraclass correlation coefficient for category gist was likely due to large within-subject changes in that index over time. Because most of the models are above threshold, we consider there to be significant individual differences in the memory variables we measured. We therefore proceeded with multi-level modelling as planned.

### Gist Memory over Time

There was no decline in inferential gist between Pre-Sleep to Post-Sleep (*p* = .739) or between Pre-Sleep and Day7 (*p* = .662). There was no significant decline from Pre-Sleep to Post-Sleep for summary gist (*p* = .068), but a significant decline from Pre-Sleep to Day7 (*p* = .000). There were significant declines across both time points for both statistical learning (*p* = .0494 and *p* = .000) and category gist (*p* = .012 and *p* = . 000). There was no significant effect of OSPAN, *ps* > .135. A visual depiction of gist variables over time, normalized to Pre-Sleep, is given in Figure 4C. P values for all multi-level models with time predictors are given in Table 3.

**Table 3.** P-values for Multi-level Models with Time and Sleep Predictors.

| Memory Variable | Inferential | Statistical Learning | Summary gist | Category Gist |
| --- | --- | --- | --- | --- |
| Confirmatory Analyses |  |  |  |  |
| Time 1 | .739 | .049<sup>^</sup> | .068 | .012* |
| Time 2 | .662 | .000* | .000* | .000* |
| PREM | .115 | .010* | .111 | .119 |
| PREM*Time | .456, .909 | .714, .559 | .108, .108 | .186, .426 |
| PSWS | .66 | .965 | .053 | .665 |
| PSWS*Time | .276, .064 | .482, .807 | .843, .793 | .276, .874 |
| TST | .994 | .060 | 0.048<sup>^</sup> | .793 |
| TST*Time | .971, .436 | .688, .379 | .015<sup>^</sup>, .026<sup>^</sup> | .973, .845 |
| Exploratory |  |  |  |  |
| Spindles | .575 | .082 | .643 | .073 |
| Spindles*Time | .469, .979 | .915, .062 | .054, .112 | .886, .500 |
| NREM | .058 | .041<sup>^</sup> | .067 | .262 |
| NREM*Time | .859, .621 | .409, .336 | .033<sup>^</sup>, .048<sup>^</sup> | .114, .244 |
| Stage 2 | .752 | .791 | .692 | .797 |
| Stage 2* Time | .720, .928 | .513, .885 | .664, .575 | .264, .826 |

\*Significant after FDR correction ( $p < .013$ ), <sup>^</sup>Significant before FDR correction ( $p < .05$ ),
PREM = Proportion of Rapid Eye Movement Sleep, PSWS = Proportion of Slow-wave Sleep, TST = Total Sleep Time, NREM = Non-Rapid Eye Movement Sleep.

### Sleep

Participants reported an average weeknight sleep of 434 minutes (*SD* = 62 min). Average total sleep time for Pre-Sleep test was 359 minutes (*SD* = 91 min; approximately 6 hours, which is also the ‘may be appropriate’ threshold set by the National Sleep Foundation^43^). Out of the participants who completed sleep diaries, 4 indicated that they were ‘refreshed’ the morning of the Pre-Sleep test, 28 indicated they were ‘somewhat refreshed’, and 26 indicated they were ‘fatigued’. However two sample t-tests between participants who indicated that they were either ‘refreshed’ or ‘somewhat refreshed’ and those who indicated that they were ‘fatigued’ did not significantly differ on any of our four gist memory measures at Post-Sleep (*p* = .551 for inferential gist, *p* = .819 for summary gist, *p* = .252 for statistical learning, and *p* = .800 for category gist). Therefore, although sleep was shorter than a recommended typical night, subjective fatigue did not appear to impair gist memory performance. Along these lines, we note that even naps have been shown to have memory effects,^38^ so shorter sleep durations should not preclude observation of possible memory effects.

Regarding our confirmatory analyses, after controlling for FDR (significance threshold of 0.05 divided by 4 variables is 0.013), proportion of REM negatively predicted statistical learning collapsed over time, or controlling for time, *b* = −0.626, *t*(58) = −2.662, *p* = .010. The role of REM on gist variables is given in Figure 3C-F. All of the following results were only significant before controlling for FDR. TST negatively predicted summary gist at Post-Sleep (*b* = −0.343, *t*(109) = −2.480, *p* = .015) and Day7 (*b* = −0.386, *t*(109) = −2.252, *p* = .026). Regarding our exploratory analyses, NREM positively predicted statistical learning over time or controlling for time, *v* = .445, *t*(62) = 2.089, *p* = .041. NREM also positively predicted summary gist at Post-Sleep, *b* = 3.325, *t*(120) = 2.156, *p* = .033, and Day7, *b* = 3.843, *t*(120) = 1.999, *p* = .048. We did not find any significant Spindle or Stage 2 effects, *ps >* .062. For the purposes of future hypotheses, we note a positive trend toward SWS positively predicting summary gist, *b* = 1.625, *t*(56) = 1.980, *p* = .053. P values for all multi-level models with time and sleep predictors are given in Table 3.

### Results of hippocampal analysis

Average anterior hippocampal volumes were 1027 mm³ (*SD* = 158 mm³) on the left and 1100 mm³ (*SD* = 123 mm³) on the right. Average posterior hippocampal volumes were 1053 mm³ (*SD* = 119 mm³) for the left, and 1026 mm³ (*SD* = 120 mm³) on the right. Average anterior/posterior ratios were 0.98 (*SD* = 0.19) on the left, and 1.09 (*SD* = 0.22) on the right. In this study, we collected data from a narrow age range (22-35) among healthy adults. We pre-registered that if age significantly predicts either anterior and posterior hippocampal volumes in a regression, we will include age as a covariate. If age does not significantly predict either or pHPC volume, then to simplify our analyses, we will not include age. Age did not significantly predict hippocampal volumes in our sample, so we did not include age in our models (*p* = .425 and *p =* .885 for left and right aHPC volumes, *p* = .084 and *p* = .084 for pHPC volumes). After controlling for FDR, we did not find any significant effects of aHPC volumes, pHPC volumes or aHPC/pHPC ratio when including these variables in our model, all *p*s > 0.09. We do note a trend for left anterior hippocampal volumes predicting statistical learning, *b* = .000, *t*(120) = 1.738, *p* = .085 and negatively predicting associative gist at Post-Sleep, *b* = -.001, *t*(115) = −1.697, *p* = .093. We also note a trend for the right anterior-posterior ratio predicting inferences at Day7, *b*= -.137, *t*(103) = −1.610, *p* = .110.

## Discussion

The goal of our study was to investigate the relationship of time and sleep stages on various forms of gist memory. We found a clear behavioural dissociation between inferential gist on the one hand, which was sustained over the course of a week, and statistical, category and summary gist on the other, which decayed over the same period. Thus, not all gist memory behaves the same over time, and among the forms of gist we evaluated, inferential gist appeared to be uniquely protected by consolidation processes supportive of long-term retention. This result is consistent with previous studies comparing sleep and wake groups.^4, 45^ Another interpretation for this result could be that inferential gist is learned quickly and does not fade over time. We also found that proportion of REM sleep negatively predicted statistical learning over time, consistent with the proposal that REM sleep may contribute to discretization of information (and thereby working against the extraction of regularities from experienced events). By contrast, hippocampal volumes were not a reliable predictor of longitudinal effects.

Taking Lewis & Durrant’s model of overlapping memory representations as a general gist extraction process,^11^ we conclude that memory overlap is an insufficient condition for long-term generalization. In our study, only structured relational memories (inferential gist), where each item was related to each other item, were enhanced. This type of memory contrasts with category memories (category gist), summary gist, and repeated co-occurrence in statistical learning, which could involve discrete sets of overlapping memories organized into triplets. Inferential gist likely involves the formation of an overarching structural or schematic framework, such that any inference or premise pair can be referenced to this framework to determine the correct response. This may be a more similar structure to insight tasks^46^ and artificial grammars.^38^ Such a framework could be revealed through the synthesis of episodes characterizing individual premise pairs.

Our finding that inferential gist is retained over sleep and time is consistent with previous research comparing sleep and wake groups.^4, 45^ Although it is possible that the retention in memory is due to repeated testing, we did not find a significant test-retest effect when testing twice during the Pre-Sleep session. Furthermore, each task was presented to participants in a counterbalanced order, and no one task was deemed more important, so it is unlikely that participants selectively encoded information from TI above other tasks. In our data, we did not find any significant predictive effects of sleep stages on inferential gist at specific time points or collapsed over time. Speculatively, the absence of interfering stimuli during sleep could be what promotes higher inferential gist memory in sleep rather than wake groups.

As in any single experiment, is possible our results were idiosyncratic to our specific choice of tasks. In future work, it will be valuable to test the generalizability of our findings to different tasks that putatively address the same constructs.

### Sleep Stages

Proportion of REM predicted reduced statistical learning controlling for time, an effect not found with total sleep time. As this effect was found collapsing over time rather than interaction with a specific time point, one possible explanation is that a waking mechanism analogous to REM sleep exists that inhibits statistical learning at initial test and other time points. As mentioned above, Landmann et al. propose that REM sleep is responsible for a process called schema disintegration, which disbands existing schemas and allows for creativity.^7^ Considering this hypothesis, participants with higher proportions of REM sleep may have weaker statistical learning schemas. In contrast to previous research, we did not find any evidence that SWS predicts inferential gist, statistical learning, summary gist, or category gist (transitive inference^45^ and statistical learning^5^). We also did not find evidence to support the theory proposed by Landmann et al. that schema integration (of which inferential gist is a special case) and schema formation (statistical learning) takes place during SWS or NREM sleep.^7^ We do note, however, a positive trend towards NREM sleep predicting statistical learning (significant before FDR correction). Hence, an active sleep mechanism to retain statistical learning may exist in addition to a negative (REM) inhibitory mechanism.

We also note a positive trend towards TST negatively predicting summary gist and NREM positively predicting summary gist at Post-Sleep and Day7 (significant before FDR correction), suggesting that while more sleep may lead to worse summary gist processing, there may be an active NREM mechanism to retain semantic gist from lists.

### Relationships among Gist and Detail Memory Measures

Looking at the relationships among gist and detail measures, we found within-task correlations across gist and detail memory in all of our tasks. Hence, it is possible that gist memory may draw upon the same underlying representation as detail memories, or alternately be inter-dependent with them: for instance, initial retrieval of a gist memory might help to align cortical state closer to that observed at encoding, thereby facilitating reinstatement of a detailed memory (see Conway & Pleydell-Pearce,^47^ for similar notions of hierarchical memory search). We also found correlations between premise pair memory, associative memory measures, and summary gist, suggesting an associative cognitive component involved in learning premise pair relationships (two shapes in TI, words and scenes in AM). This idea is also consistent with the lack of a correlation between premise pairs and detail memory, as during detail questions participants had to distinguish only the correct detail arrangement they had seen, and did not need to retrieve the word-scene association. Lastly, summary gist memory was correlated with statistical learning and associative memory, which were not correlated with one another. Summary gist has a gist extraction component, as well as a temporal component as during study, individual words move quickly on the screen. VSL required participants to watch a quickly moving sequences of shapes and consolidate them immediate into patterns, which may be the cognitive component correlated to summary gist.

Brainerd and Reyna^48^ suggested that gist traces are more resistant to forgetting than detail traces. Our omnibus MANOVA for AM and DRM tasks was significant, with follow-up test indicating significant decreases only in associative gist and instance memory. Hence, associative gist traces are more rapidly forgotten over time than detail traces and detail may be preferentially consolidated in this task. This goes against our earlier hypothesis that gist memory would be more resistant to forgetting than detail memory. Our omnibus MANOVA for transitive inference was not significant, so we did not find any evidence that inferential gist memory (TI) is more resistant to decay than premise pairs (TI).

### Hippocampal Volumes

We did not find any significant hippocampal effects. Previous studies have found activation in the right aHPC during overlapping transitive inference pairs,^49^ as well activation in the aHPC in more distant (compared to more proximal) premise pairs.^50^ Thus, although inferential gist is likely to be encoded in the aHPC, larger aHPC volumes do not seem to predict better inferential gist memory. Previous research also found activation in the right hippocampus during VSL.^51^ One obvious distinction between these studies and ours is that we measured individual differences in hippocampal volumes, rather than activation in the aHPC and pHPC, and there is rarely correspondence across these measures. However, we did observe trend-level relations that left anterior hippocampal volumes positively predicted statistical learning, but negatively predicted associative gist at Post-Sleep, which may warrant investigation in future research.

The current design did not allow us to pinpoint the dynamic changes in hippocampal activation and sleep over time. Although we were able to behaviorally measure different kind of gist and relate those to individual differences in hippocampal volumes, there may be time-dependent changes in the activation of the aHPC and pHPC. For instance, in a recent paper, Dandolo and Schwabe^28^ found that activity in the aHPC significantly decreased over time and was related to memory specificity at encoding, whereas pHPC activity remained the same. Tompary and Davachi^52^ found that feature overlaps emerged in the hippocampus over the course of a week. Future studies could test within an fMRI over three sessions to relate activation in the aHPC to pHPC to time-dependent memory changes.

### Conclusion

Because only structured relational memories, rather than associative memories (category gist) or repeated co-occurrence (statistical learning) were retained over the course of a week, we conclude that inferential gist follows a different mnemonic trajectory than other forms of gist. Furthermore, REM sleep may be involved in schema disintegration, which although potentially beneficial for distinguishing similar events, works against participants’ ability to identify contiguous series.

## Acknowledgements

This research was funded by Natural Sciences & Engineering Research Council Discovery Grant 03637 (J.P.), which also supported N.M. Infrastructure funding was provided by Canada Foundation for Innovation – John R. Evans Leaders Fund (J.P.), and a Queen’s University Research Initiation Grant to J.P., who was supported by the Canada Research Chairs program. We gratefully acknowledge Gillian Marvel for assistance with behavioural and sleep EEG data collection; Julie Tseng, Lauren DeMone, and Natalie Doan with scheduling; Julie Tseng and Don Brien with MRI data acquisition; and Justin Siu, Roland Dupras and Mike Lewis with technical support; Holly Mcdougall for modelling and Eric Brousseau for photography.

## Disclosure Statement

Financial Disclosure: none.

Non-financial Disclosure: none.

A pre-print of this manuscript is available on bioRxiv.

